# When a phenotype is not the genotype: Implications of phenotype misclassification and pedigree errors in genomics-assisted breeding of sweetpotato [*Ipomoea batatas* (L.) Lam.]

**DOI:** 10.1101/747469

**Authors:** Dorcus C. Gemenet, Bert De Boeck, Guilherme Da Silva Pereira, Mercy N. Kitavi, Reuben T. Ssali, Obaiya Utoblo, Jolien Swanckaert, Edward Carey, Wolfgang Gruneberg, Benard Yada, Craig Yencho, Robert O. M. Mwanga

## Abstract

Experimental error, especially through genotype misclassification and pedigree errors, negatively affects breeding decisions by creating ‘noise’ that compounds the genetic signals for selection. Unlike genotype-by-environment interactions, for which different methods have been proposed to address, the effect of ‘noise’ due to pedigree errors and misclassification has not received much attention in most crops. We used two case studies in sweetpotato, based on data from the International Potato Center’s breeding program to estimate the level of phenotype misclassification and pedigree error and to demonstrate the consequences of such errors when combining phenotypes with the respective genotypes. In the first case study, 27.7% phenotype misclassification was observed when moving genotypes from a diversity panel through *in-vitro*, screenhouse and field trialing. Additionally, 22.7% pedigree error was observed from misclassification between and within families. The second case study involving multi-environment testing of a full-sib population and quantitative trait loci (QTL) mapping showed reduced genetic correlations among pairs of environments in mega-environments with higher phenotype misclassification errors when compared to the mega-environments with lower phenotype misclassification errors. Additionally, no QTL could be identified in the low genetic correlation mega-environments. Simulation analysis indicated that phenotype misclassification was more detrimental to QTL detection when compared to missingness in data. The current information is important to inform current and future breeding activities involving genomic-assisted breeding decisions in sweetpotato, and to facilitate putting in place improved workflows that minimize phenotype misclassification and pedigree errors.

## Introduction

It is a generally accepted concept that the environment in which an organism is placed affects the expression and function of genes responsible for a trait (Allard and Bradshaw, 1964; Baye et al., 2011). The magnitude of phenotypic plasticity to adapt to different environments is genotype-dependent, hence the environment can interact with a genotype to shape the phenotypic traits, leading to genotype-by-environment (GE) interaction (Genard et al., 2017). In plant breeding, GE interaction is expressed as either genotypic rank-change among genotypes due to varied responses to changing environments or as absolute change in trait values without a rank change (Crossa, 2012). Since these interactions are unpredictable as the environments themselves, they confound breeding efficiency and reduce genetic gains from plant breeding (Crossa, 2012; Osei et al., 2018).

The need to account for GE interactions in making plant breeding decisions has become even dire with the current advent in applying genomic selection to increase breeding efficiency. Defined by Meuwissen et al. (2001), genomic selection is a breeding tool that uses information from all molecular markers across the genome to predict the breeding value of an individual. To be applied, this tool requires testing of models using phenotypic and genotypic information from a sample of the breeding population selected to represent the diversity (training population) in the said breeding population that is targeted for prediction (prediction population). This approach therefore calls for generating both phenotypic and genotypic data of the training population, and only genotypic data for the untested prediction population. The development of next-generation, high-throughput genotyping methods like genotyping-by-sequencing (Elshire et al., 2011) have drastically reduced genotyping costs thereby enhancing generation of large volumes of genotypic data quite fast. This has therefore left phenotyping as the bottleneck in plant breeding.

Precise phenotypic data of the training population is a prerequisite for improving the accuracy of predicting untested genotypes in genomic selection models (Velazco et al., 2017). However, GE interactions are known to increase with increasing number of genotypes and environments. Allard and Bradshaw (1964) showed that GE interactions calculated as 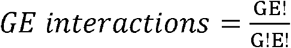, lead to exponential increase in interactions as both genotypes and environments increase. For example, they showed that two genotypes in two environments would result in about four GE interactions while 10 genotypes in 10 environments would result in up to 400 GE interactions. Plant breeding experiments always deal with far greater numbers of genotypes. Additionally, the unbalanced nature of the number of genotypes and experimental designs in most plant breeding experiments increases heterogeneity thereby complicating the variance-covariance structures of phenotypic observations (Bernal-Vasquez et al., 2014). Linear mixed models have been generally applied to analyze for GE interactions in plant breeding experiments (Piepho, 1997; 1998; Piepho and Moehring, 2005; Smith et al., 2005, Crossa et al., 2006).

The sheer large number of genotypes tested in early breeding stages means that experimental plots are large hence leading to local heterogeneity within experiments. To further improve prediction accuracies, different spatial adjustment models have been fronted to help deal with heterogeneity in experiments especially in these large early stage breeding trials (Lado et al., 2013; Bernal-Vasquez et al., 2014; Piepho et al., 2015; Velazco et al., 2017; Ward et al., 2019). Multidisciplinary teams are therefore continually working to improve the precision of measuring the phenotypes of the training populations to improve predictive ability of genomic selection in plant breeding, as summarized by Ward et al. (2019). Several of these teams have shown that considering GE interactions and spatial adjustments contributed to increased predictive ability. Lado et al. (2013) showed increased predictive ability with spatial adjustment of trial data in wheat. Elias et al. (2018) showed increase in predictive ability by about 3.4% in cassava following spatial adjustment. Ward et al. (2019) showed that correcting for spatial variation improved across location heritability by 25% but not prediction accuracy whereas correcting for GE interactions increased prediction accuracy by 9.8% in early breeding stage evaluation of wheat.

Whereas random GE interactions and spatial variation have been statistically proven to affect the precision of measuring the phenotype, the question that is not often answered is: how much of the variation observed from one experiment to the next is actually due to GE interactions? Although already a known problem in the statistical world (Schlimmer and Granger, 1986), with suggestions on data quality and cleaning (Rahm and Do, 2000; Guillet and Hamilton, 2007), experimental noise especially resulting from human error is the most difficult to correct using statistical methods. Such errors are mainly due to mislabeling, hence misclassification of study genotypes in different experiments which may also be presented as GE interaction in data. Despite this, there are currently very few studies addressing and reporting experimental noise in plants (Biscarini et al., 2016).

Sweetpotato is an important crop for food and nutrition security especially in sub-Saharan Africa (SSA). Having a complex, autohexaploid, genome ensured that genomics-assisted breeding has lagged behind for this crop. However, global efforts are now in place to ensure that new breeding tools such as genomic selection and marker-assisted selection are applied to benefit small-holder farmers and consumers of sweetpotato in SSA. In the current study we aimed to answer the following questions: i) how much mislabeling can be expected through different stages of population development for trialing within a breeding program, ii) what would be the effect of such mislabeling mistakes on marker-trait associations, iii) what are the effects of different proportions of mislabeling versus missingness using simulations on real data; iv) what would be the implications of such findings on designing a genomics-assisted breeding strategy for sweetpotato. We use two case studies and simulation based on data from some of the sweetpotato populations being used for genetic studies in preparation for deploying genomics-assisted breeding methods for sweetpotato improvement. All data are based on the global sweetpotato breeding program of the International Potato Center (CIP) through its various regional and sub-regional breeding platforms.

## Materials and Methods

### Case Study 1: Genetic fidelity in the Mwanga Diversity Panel (MDP), a genetic study breeding population

#### Genetic materials

The MDP population was developed from the sweetpotato breeding platform for east and central Africa of the International Potato Center (CIP). It is made up of a diallel cross among 16 parents from this breeding platform, coming from two gene pools (here A and B) separated by SSR markers (David et al., 2018). There are 64 families (8B by 8A crosses) each with 30 genotypes on average. Sweetpotato is mainly outcrossing, self-incompatible and heterozygous. Apart from the crossing step where propagation is by seed, sweetpotato is clonally propagated throughout the other stages of the breeding process. Therefore, each seed is potentially a different genotype. This population was established for the purpose of genetic studies in developing tools for genetic linkage in multi-family breeding populations, genome-wide association mapping and genomic selection for complex autopolyploid genomes

#### Population establishment and trialing

The process of establishing this population is shown in the flowchart below (Figure 1). In summary, the crossing among parents was done in CIP-Uganda, where the seed inventory was established. Then seed was shipped to CIP-Kenya for *in vitro* germination, where the population is maintained *in vitro*. Sweetpotato, which almost behaves like a weed, is not *in vitro-*friendly hence requires constant multiplication and regeneration *in vitro*. Also, sweetpotato seed requires scarification protocols to germinate the seed which means that not all seed were successfully germinated the first time per family and new seed shipments were required for such families from CIP-Uganda to CIP-Kenya. After *in vitro* establishment, the population was then shipped back to CIP-Uganda for trialing. Since the population could not be established *in vitro* at the same time, *in vitro* plantlets were also shipped back to CIP-Uganda from CIP-Kenya in batches.

**Figure 1.**
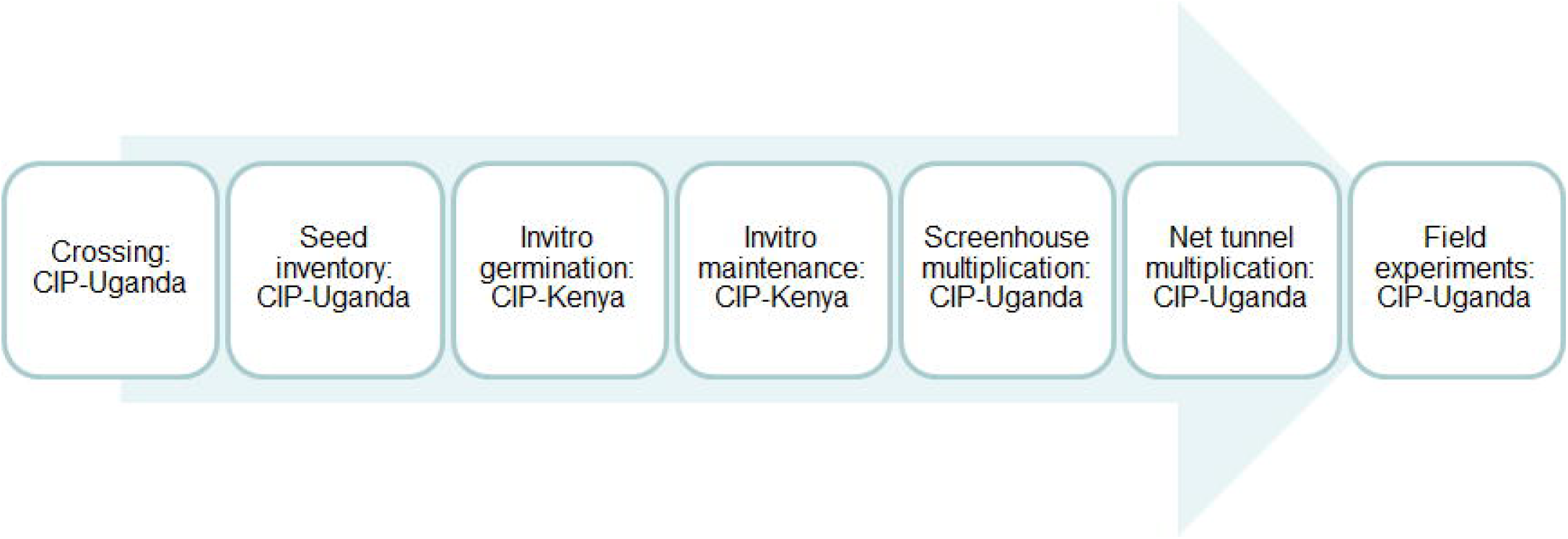
Flow chart of population development of the Mwanga Diversity Panel (MDP) population from crossing to field evaluation

As each individual seed in sweetpotato is a potential new variety, the *in vitro* plantlets need to be grown in a screenhouse for cloning. Additionally, for experimentation, screenhouse plants need to be multiplied further through vines to have enough planting materials for experiments. The most important virus disease for sweetpotato is sweetpotato virus disease (SPVD), a complex caused by the synergistic interaction of *Sweet potato feathery mottle virus* and *Sweet potato chlorotic stunt virus*, transmitted by aphids and whiteflies, respectively (Clark et al., 2012). Since the CIP support platform in Uganda is located in Namulonge, a hotspot for SPVD, the vines cloned from screenhouses need to be multiplied in net tunnels that keep away the virus vectors to furnish virus-free planting materials for experiments. As part of quality control (QC) and quality assurance (QA), each net tunnel is planted with only genotypes from the same family. The vines for experimentation are taken from these net tunnels to the field experiments in Uganda. As part of QC/QA also, the two teams at CIP-Uganda and CIP-Kenya worked closely together. However, with a large population, barcodes were not used at all stages hence anticipation of some degree of human error.

#### Molecular quality control for genetic fidelity through the various transfer stages

During the first season of field trialing (2018), we randomly sampled about 5% of the population from *in vitro*, screenhouse and field experiments from one of the three field locations. Given that this is a breeding population developed from a diallel cross, our specific objectives were: i) to evaluate genetic fidelity following movement from *in vitro* to screenhouse and then to field; ii) to examine the level of mislabeling between and within families due to the population establishment process. The field and screenhouse sampling were done at the National Crop Resources Research Institute (NaCRRI) in Uganda, while *in vitro* sampling was carried out in the Biosciences eastern and central Africa - International Livestock Research Institute (BecA-ILRI) Hub based in Nairobi, Kenya. The random sampling resulted in 13 out of the 64 families sampled, with an average of seven genotypes per family resulting in 94 samples hence three DNA plates were sent for genotyping (one each from *in vitro*, screen house and field). Although structure analysis does not require a high-density marker set, the complexity of the sweetpotato genome has ensured that a QC/QA low density marker set for routine use was still unavailable for sweetpotato breeding programs at CIP and African National Research Institutions (NARIs). Therefore, the three plates were genotyped at high density using Diversity Arrays Technology’s (DArT) sequencing-based technology (DArTseq) implemented by the Integrated Genotyping Service and Support (IGSS), based at BecA-ILRI in Nairobi. The high-density genotyping resulted in about 41,194 SNPs (**Supplementary Table 1**). Hard filtering of these based on polymorphic information content (PIC) ≥ 0.25, minimum allele frequency ≥ 20% and call rate ≥ 90% left 11,622 SNPs that were used to develop a distance matrix and a phylogenetic tree. We used diploidized SNPs (SNPS without ploidy dosage information) for this study. The distance matrix and phylogenetic tree were generated using DARwin 6.0.21 (Perrier and Jacquemoud-Collet, 2006). Afterwards the clustering was examined based on positions on the tree and Sankey diagrams developed using the Alluvial package (Bojanowski and Edwards, 2016) in R.

## Results of the QC experiment

The phylogenetic tree based on distance matrix (Figure 2) indicated expected clustering of same genotypes from *in vitro*, screenhouse and field of a larger percentage of the genotypes tested. However, an additional percentage of tested genotypes such as I28 (meaning *in vitro* 28) and its counterparts in the screenhouse and field, S28 and F28, respectively, did not cluster as expected indicating a level of mislabeling error **(Supplementary Figure 1)**. The summary of the tree order, genotype names, phenotypically assigned families, families suggested by the genetic distance matrix, and the female and male parents of each phenotypically assigned family are shown in **Supplementary Table 2**. Analysis of the tree order indicated that 26 out of 94 genotypes tested did not cluster as expected among *in vitro*, screen house and field samples thereby indicating about 27.7% labeling errors as the germplasm moved from *in vitro* to screenhouse and then to the field (Figure 3, **Top**). Analysis for between and within family mislabeling indicated that 64 out of the 282 tested genotypes did not belong to the phenotypic assigned families as indicated by genetic distance. This indicated that we had about 22.7% mislabeling error between and within families (Figure 3, **Bottom**).

**Figure 2.**
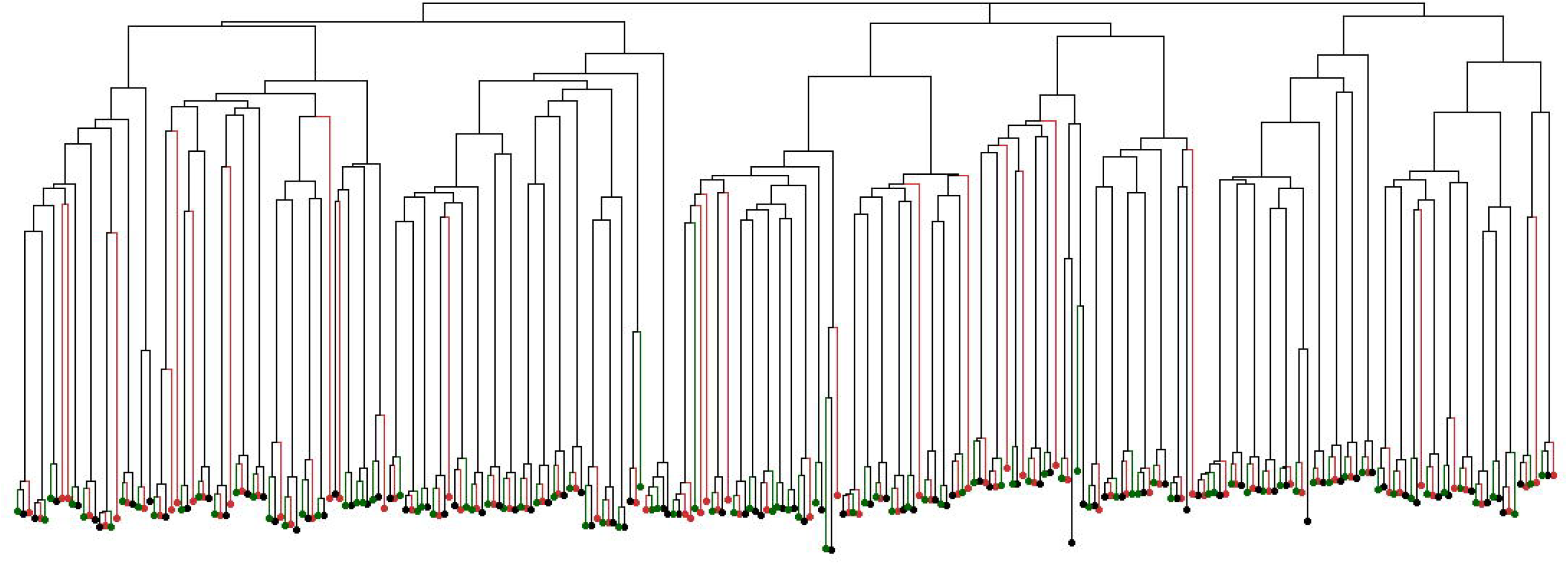
Unweighted neighbor-joining tree based on dissimilarities of 94 genotypes replicated from field, screenhouse and *in vitro* (hence 282 samples in total) using 11,622 DArTseq SNP markers. Green dots indicate samples from field, black dots indicate samples from screenhouse, and red dots indicate samples from *in vitro*.

**Figure 3.**
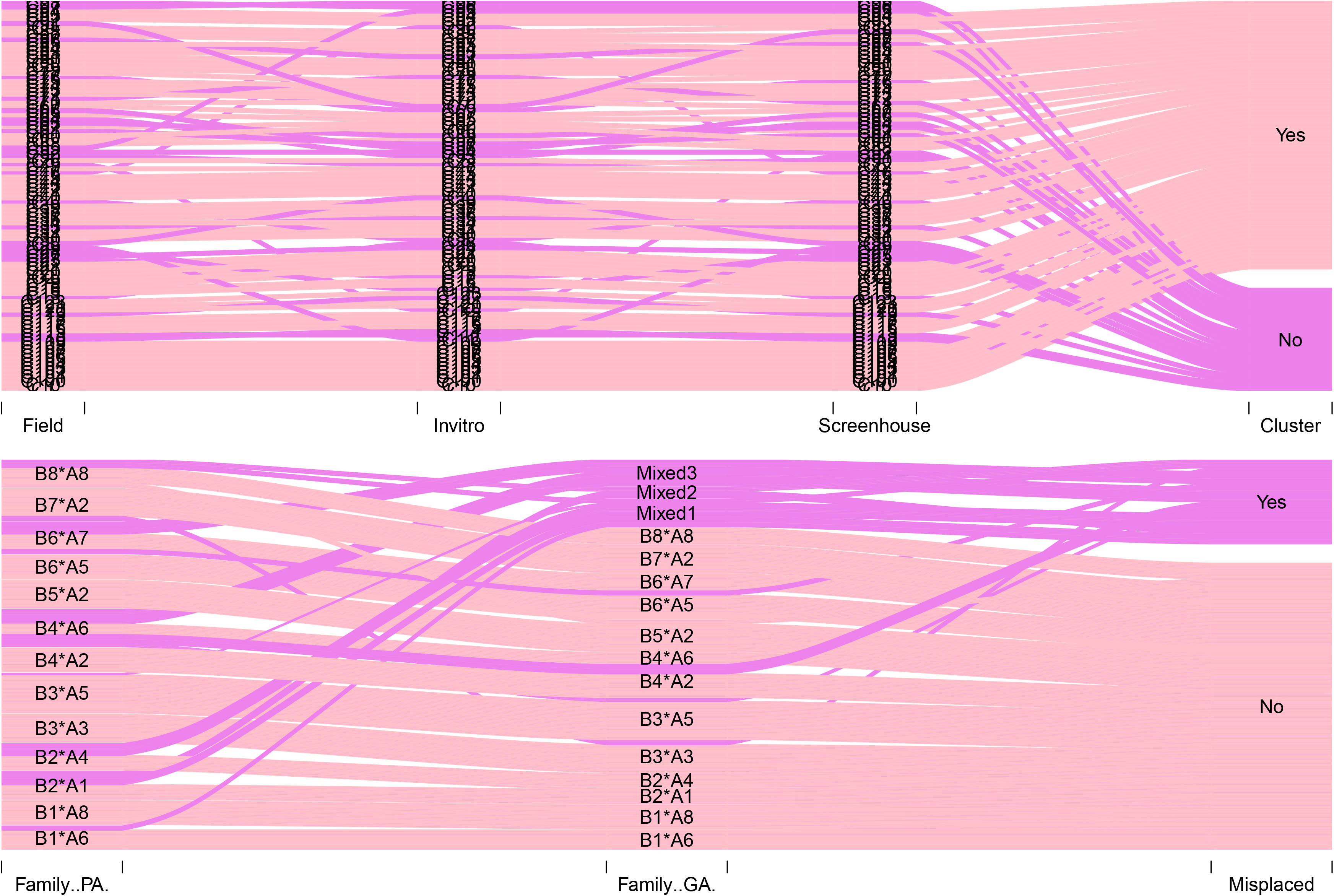
Top: A Sankey diagram showing mislabeling from *in vitro*, screenhouse and field of 94 randomly selected genotypes of the MDP population based on the tree order of the genetic distance matrix using 11,622 SNP markers. The y-axis indicates the cluster order of the genotypes from the phylogenetic tree. The purple color indicates those that are mislabeled while the pink color indicates genotypes which cluster well from *in vitro*, screenhouse and field. Bottom: A Sankey diagram showing mislabeling among different phenotypically assigned families (Family PA) and clustering based on genetic distance matrix (Family GA). The mixed families Mixed1, Mixed2 and Mixed3 were assigned when more than two families appeared on the same clade of the phylogenetic tree. The y-axis represents the family names. The purple color represents misplacement within families, the pink color represents agreement in family assignment between phenotypic and genetic distance.

### Case Study 2. Quantitative trait loci-by-environment (QTL x E) analysis of a biparental population

#### Genetic materials

A 315-progeny biparental population developed from a cross between Beauregard and Tanzania cultivars was evaluated in a multi-environment testing (MET) experiment in three countries: Peru, Ghana and Uganda. The two parents segregate for various traits of interest in sweetpotato such as β-carotene, starch, dry matter and yield related traits. Beauregard is a US-bred variety while Tanzania is an African farmer selected variety. Additional information about this population can be found in Pereira et al. (2019) and Gemenet et al. (2019**; submitted)**.

#### Population establishment and trialing

The chain of trial establishment is presented in Figure 4. Crossing of the two parents, seed inventory, *in vitro* germination/maintenance, DNA extraction were all carried out in CIP-Peru. DNA was then shipped to the Genomic Science Laboratory (GSL) at the North Carolina State University (NCSU) for genotyping. Additionally, the *in vitro* plantlets were shipped to Ghana and Uganda. Screenhouse/net tunnel and field multiplication for trialing was carried out in Peru, Ghana and Uganda. After multiplication, six field experiments were carried out in three locations of Peru over two years (2016-2017), eight field experiments were carried out in three locations of Ghana over three years (2016-2018), and six field experiments were carried out in three locations of Uganda over two years (2017-2018). The trials in Peru were grown in Ica (latitude 14° 01′ 44.7″ S, longitude 75° 44′ 37.5″ W), San Ramon (11°07′29″S, 75° 21′ 25″ W) and Pucallpa (8° 23′ 34.3″ S, 74° 34′ 57.4″ W). In Uganda, the experiments were grown in Namulonge (0° 31′17.99″ N, 32° 36′ 32.39″ E), Serere (1° 29′ 59.99″ N, 33° 32′ 59.99″ E) and Kachwekano (1° 15′ 0″ S, 29° 57′ 0″ E). In Ghana, the experiments were grown in Wenchi (7° 44′ 0″ N, 2° 6′ 0″ W), Fumesua (6° 42′ 39.41″N, 1° 31′ 2.03″W) and Nyankpala (9° 24′ 0″ N, 0° 58′ 60″ W). In Peru, the experiments in Ica were grown under two treatments: terminal drought (where irrigation was stopped at 70 days after transplanting (DAT)) and control (optimal) where irrigation was continued until harvest at 120 DAT. They are hereby abbreviated as Ica16D, Ica16C, Ica17D, Ica17C, indicating the location, year and treatment (D = drought; C = control) while the experiments in San Ramon and Pucallpa were grown only under optimal conditions in 2016, hereby abbreviated as SR16 and Puc16, respectively. In Uganda, all experiments were grown under optimal conditions and abbreviated as Nam16 and Nam17 for Namulonge in 2016 and 2017 respectively, Ser16 and Ser17 for Serere in 2016 and 2017 respectively, and Kac16 and Kac17 for Kachwekano in 2016 and 2017 respectively. In Ghana, except for Nyankpala and Fumesua in 2016 both abbreviated as (Nya16 and Fum16, respectively), all other experiments were grown under terminal drought and control (optimal) treatments as described for Peru. They are abbreviated as Wen17D and Wen17C for Wenchi; Nya17D, Nya17C, Nya18D and Nya18C for Nyankpala 2017 and 2018 respectively (D = drought; C = control). Locations are shown in **Supplementary Figure 2**.

**Figure 4.**
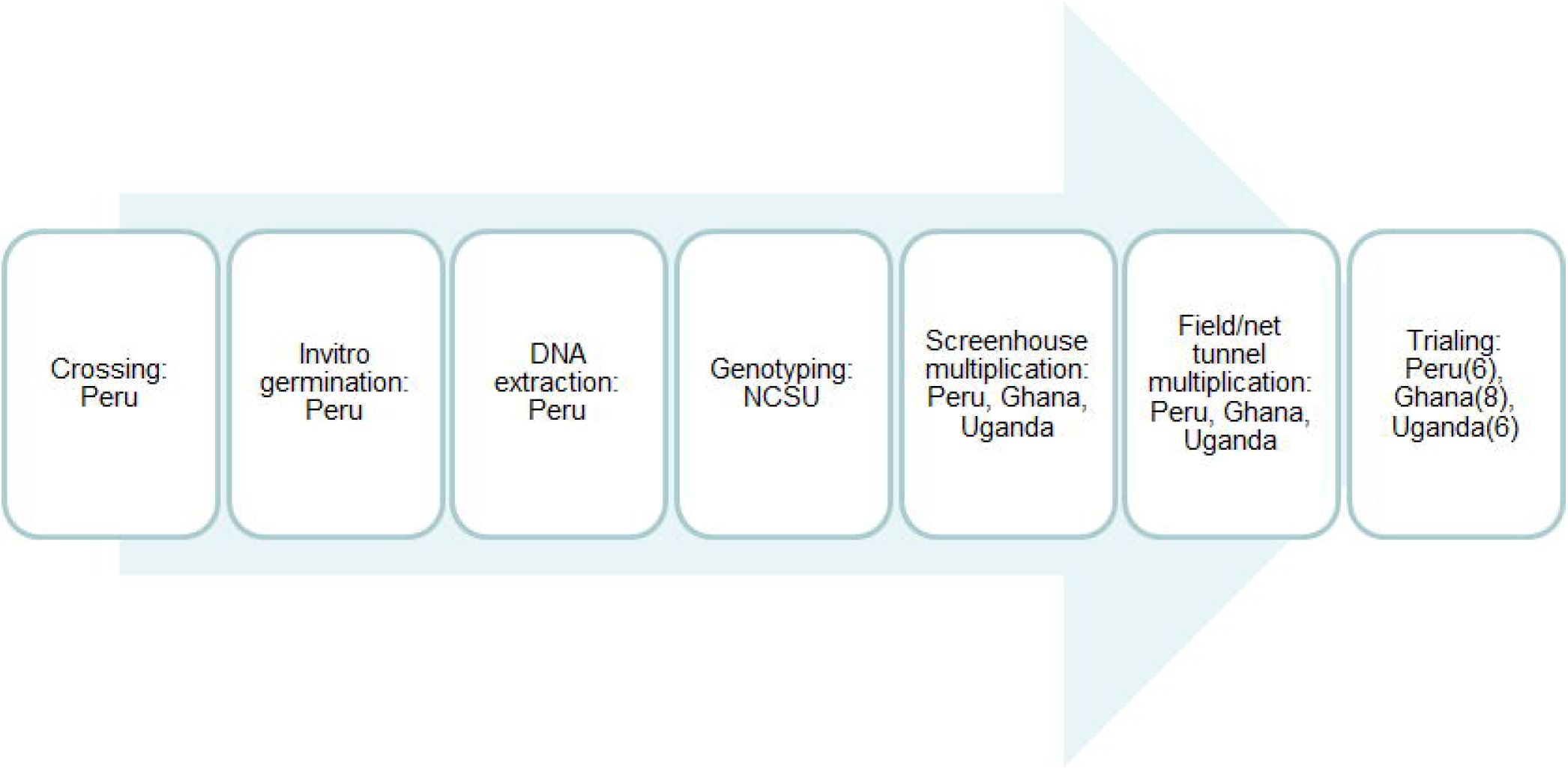
Chain of events from crossing to trial establishment in three countries with sweetpotato breeding platforms of the International Potato Center (CIP).

All the 315 genotyped progeny and parents were evaluated in Peru and Uganda. In Ghana, due to problems in multiplication, subsets ranging from 238-270 genotypes were evaluated in the eight experiments. The design was alpha lattice for all experiments in Peru, while randomized complete block design was used for experiments in Ghana and Uganda, each with at least two replications. Several yield- and quality-related traits were measured in these trials as described in Pereira et al. (2019) and Gemenet et al. (2019; submitted). Data were collected per plot and converted to per hectare based on plot sizes per experiment. For the purposes of this case study, we used only the total storage root yield in tons per hectare (rytha), for two reasons: First this trait is easier to standardize measurement across the different trials without introducing too much bias. It is measured by weighing all storage roots per plot regardless of whether they are of marketable size or not. Separating roots into marketable and non-marketable size creates subjectivity as there is not an automated way and breeders in these regions use an estimation (i.e. anything less than 100g is non-marketable and vice-versa, which is subjective as the size is by visual estimation). Secondly, because storage root yield is our primary trait in addition to other quality attributes. Our objectives were: i) to calculate genetic correlations between pairs of environments; ii) to define mega-environments among the study test sites; iii) to map QTL within and between mega-environments; iv) to simulate different proportions of misclassification through permutation, and missingness to estimate their effects on QTL detection. The raw data used in this analysis is presented in **Supplementary Table 3**.

#### Data analysis

##### Phenotypic data

To analyze the phenotypic data a two-stage multi-environment testing (MET) analysis approach was applied because different experimental designs were used across environments. In the first stage, single environment analyses were performed for all environments individually. A mixed model, taking into account the respective experimental design, was fitted to the phenotypic data (rytha trait). When plot coordinates were available, i.e. for the Peru trials, and when spatial field effects were significant, a spatial adjustment was incorporated in the mixed model. Filtering out such significant spatial field effects reduces the residual noise so that the actual genetic signal becomes more pronounced. Genotype was considered as a fixed effect in these mixed models, so that best linear unbiased estimators (BLUEs) for the genotypic rytha means were obtained per environment. In the second stage, another mixed model was fitted to the table of estimated means. A weighting scheme based on the standard errors of the estimated genotype means per environment was used in this mixed model. This meant that, on average, more weight was given to trials with a higher heritability. Based on the genetic correlations between environments estimated using this fitted mixed model, mega-environments were determined. Finally, and in a similar way, a mixed model was fitted using only estimated genotype means from the environments belonging to a mega-environment, which was then used for inference about that specific mega-environment. The genetic variances of, and the genetic correlations between, environments belonging to a certain mega-environment were estimated, and best linear unbiased predictors (BLUPs) across that mega-environment. Also BLUEs across each mega-environment were estimated by fitting a similar mixed model taking genotype as a fixed effect. The BLUEs were then used in QTL mapping.

##### Mapping of quantitative trait loci in mega-environments

Genotyping of the mapping population was done using the GBSpoly protocol optimized for sweetpotato and described by Wadl et al. (2018). Variant calling and dosage assigning for the hexaploid data was carried out as described in Pereira et al. (2019) and Mollinari et al. (2019). QTL mapping was carried out based on the phased genetic linkage map (Mollinari et al., 2019) developed using the MAPpoly program (Mollinari and Garcia, 2019) optimized for polyploids. The genetic linkage map is available interactively at https://gt4sp-genetic-map.shinyapps.io/bt_map/. The QTL analysis was done following the random effect model approach developed for polyploids and described by Pereira et al. (2019). Analysis of QTL for single environments (SE) within mega environments (ME) was carried out based on the BLUEs from the first analytical stage of the two-stage mixed model analysis described above. QTL analysis at the ME level was carried out using BLUEs from the second analytical stage.

##### Simulations to compare effects of missingness vs misclassification on QTL mapping

In order to assess the detection rate of previously identified QTL, we performed QTL analyses with increasing proportion of randomly permuted individuals, to represent misclassified individuals (200 simulations each at 10%, 20%, 30%, 40% and 50%) using rytha and flesh color (FC) adjusted means. The simulations were based on data from Peru only whose quality had been upheld and for which QTL have already been reported (Pereira et al., 2019; Gemenet et al., 2019; submitted). Subsequently, we replaced the permuted individuals from each simulation with missing data and carried out new QTL analyses on these reduced samples to represent missingness. We chose to include flesh color together with rytha in the simulation study to represent both complex and simple traits respectively. As for the MET data above, QTL detection for the simulated data was carried out using QTLpoly software (Pereira et al., 2019) based on a forward search followed by a backward elimination with the respective pointwise thresholds of p< 0.01 and 0.001.

#### Results

##### Single environment analysis

Significant genotypic variation was observed among genotypes in all single environments (SE) as indicated by box plots of BLUEs in Figure 5. It is also evident from Figure 5D that considering spatial variation significantly improved correlation among single experiments in Peru which had a field map for rows and columns to allow for some spatial adjustments, as compared to Ghana and Uganda which did not. We dropped Kac17 and Nya16 from further analysis after preliminary analysis showed that the yields of these trials were extremely outlying, relatively low and high respectively, compared to the other trials in the same country.

**Figure 5.**
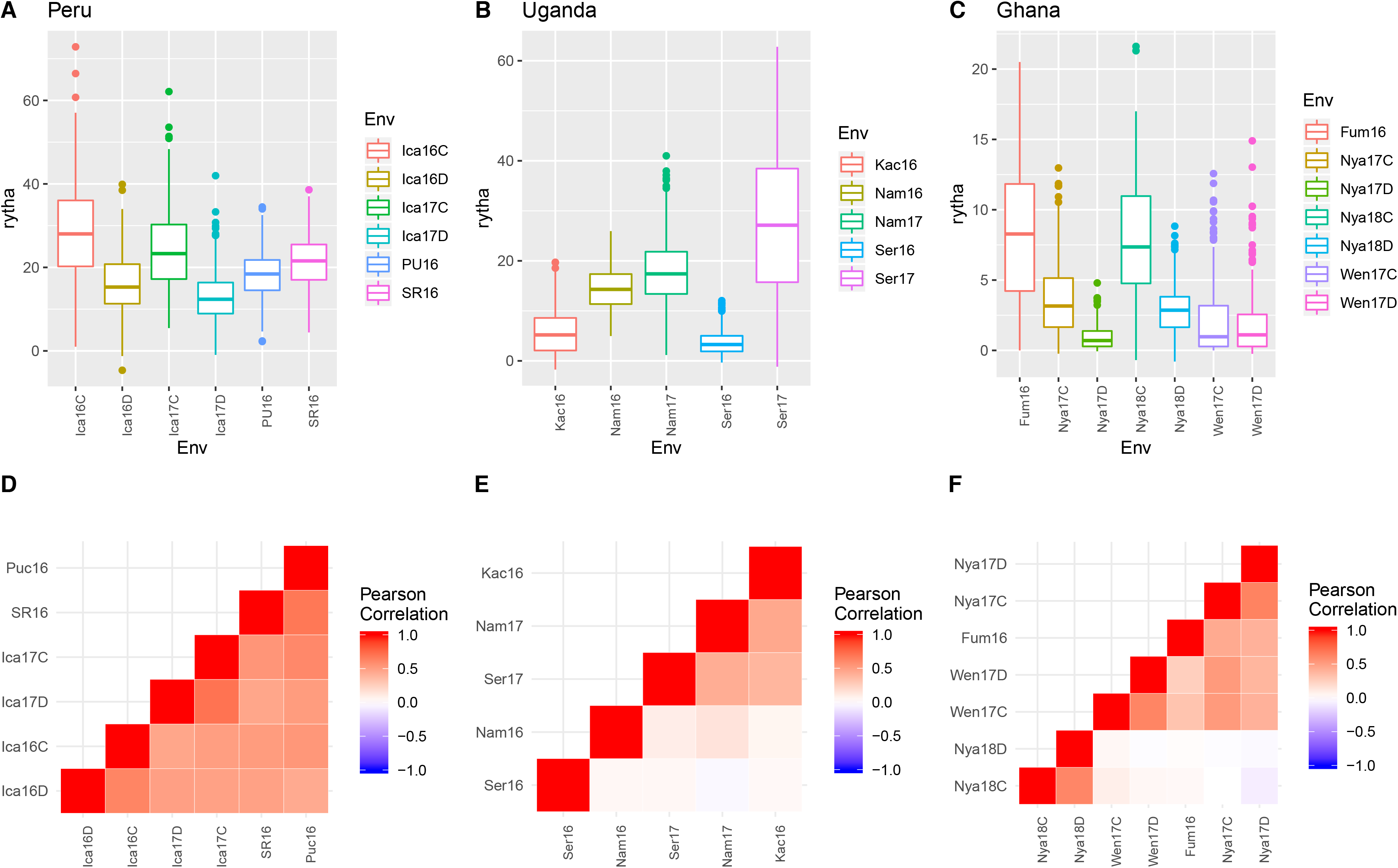
Boxplots (A, B, C) and correlation charts (D, E, F) among best linear unbiased estimators (BLUEs) for total storage root yield measured in 18 single experiments in Peru, Uganda and Ghana.

##### Multi-environment analysis

At the second stage of the MET analysis, genotype distributions among BLUPs per experiment indicated genetic variation among the 18 experiments taken together, as indicated by boxplots based on BLUPs (Figure 6A). Additionally, correlations among BLUPs for the 18 environments indicated two clear MEs (Figure 6B). Only two of the experiments in Ghana showed some correlation with some experiments from Peru and Uganda, and these were not correlated with the other experiments in Ghana. ME1 was made up of five experiments from Ghana, while ME2 was made up from 11 experiments: six from Peru, three from Uganda and two from Ghana, though the two environments from Ghana were less correlated with the rest. For further analyses, we chose to use only nine experiments from Uganda and Peru which had higher correlations among BLUPs of ME2. Genetic correlations among pairs of environments were low-to-moderate ranging from *r = 0.29* to *r = 0.65* in ME1 (five Ghana environments; Figure 6C) and moderate-to-high, ranging from *r = 0.37* to *r = 0.99* in ME2 (nine environments from Peru and Uganda; Figure 6D).

**Figure 6.**
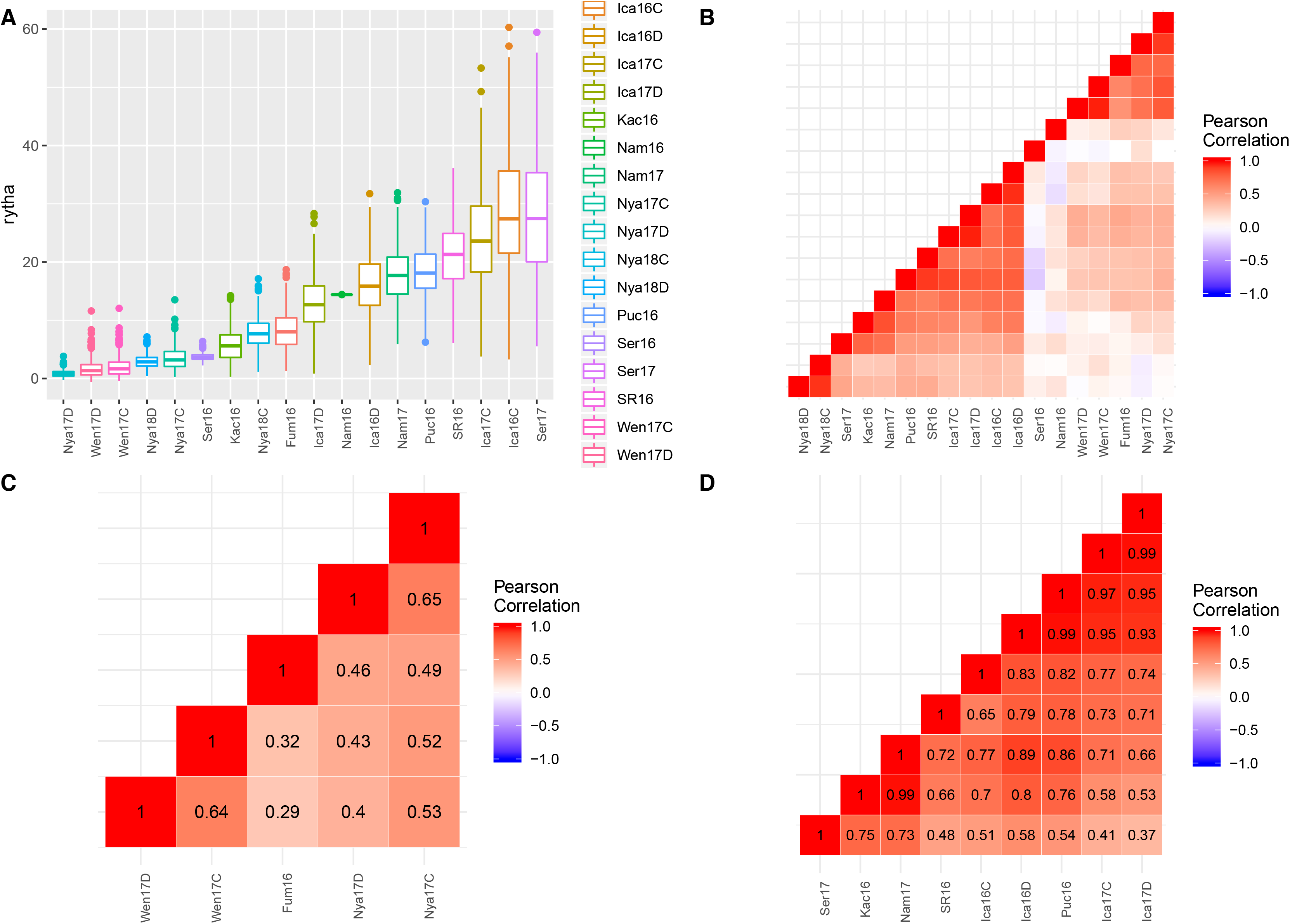
Boxplots (A), correlations and mega-environments (ME) among best linear unbiased predictors (BLUPs; B) and genetic correlations within ME1 (C) and ME2 (D) from MET analysis of 18 experiments from Peru, Uganda and Ghana.

##### Quantitative trait loci (QTL) analysis

Analyzing for QTL within the two mega-environments captured only one QTL for ME2 on linkage group (LG) 15 and no QTL for ME1 (Figure 7A). The QTL explained 16.7% of the observed variation in rytha across the ME2. Consequently, we analyzed for QTL for the single environments in ME2. Four distinct QTL were identified for SEs in ME2: one QTL was on LG 3, one on LG 13 and two on LG15 (Figure 7B; Table 1). The second QTL on LG 15 was associated with Nam16, an environment in Uganda, while the rest of the QTL were associated with environments from Peru (Table 1). Individual QTL explained between 10.9 and 22.1% of the total observed variation for rytha (Table 1). Allelic effects analysis of parental haplotypes for the ME2 QTL on LG 15 indicated that Beauregard contributed two alleles that increased and three alleles that reduced rytha whereas Tanzania contributed three alleles each to the increase and reduction of rytha respectively (Figure 7C). Results also indicate that Tanzania contributed more to the increase in rytha in ME2, when compared to Beauregard (Figure 7C).

**Figure 7.**
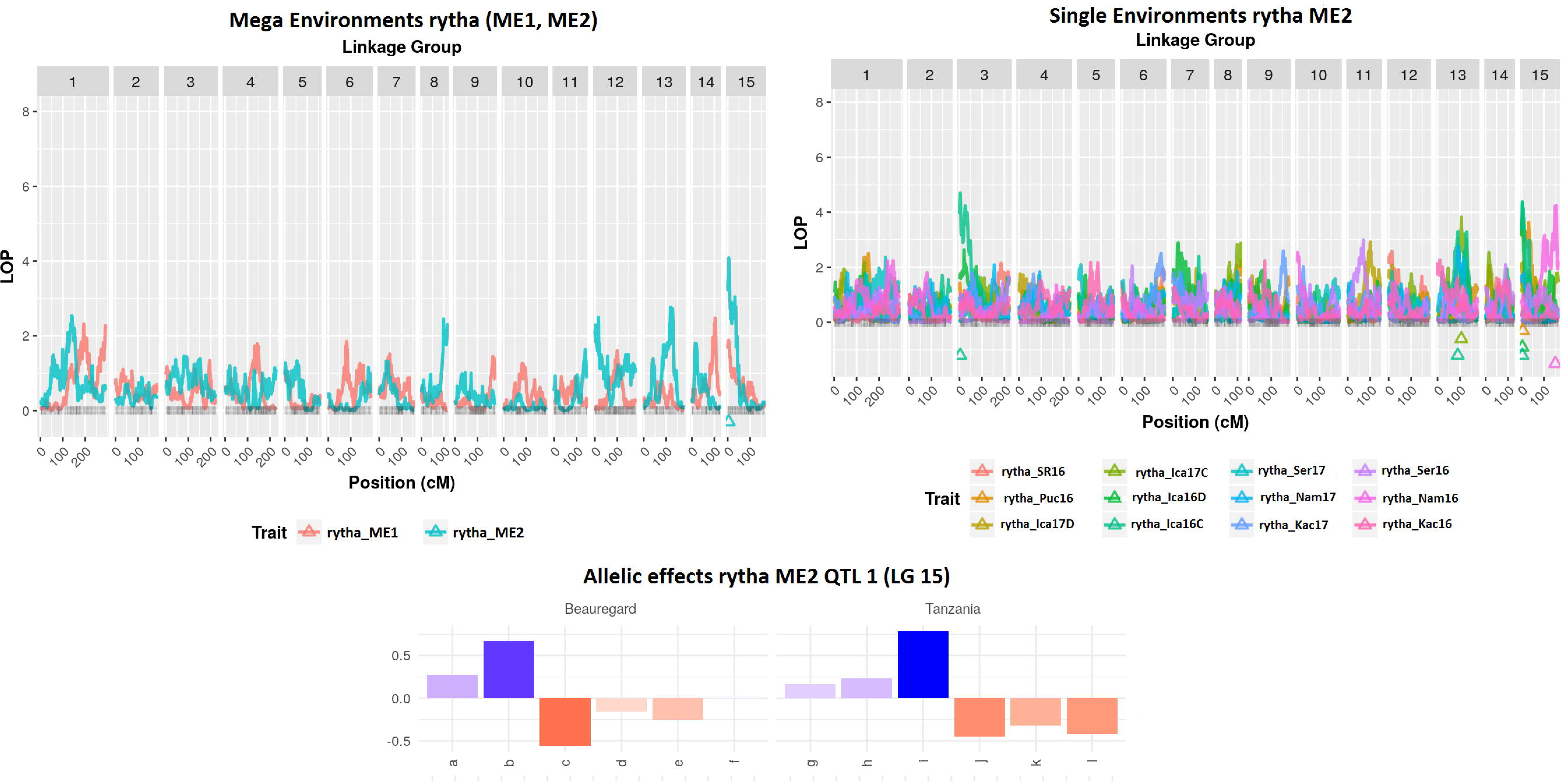
QTL plots for mega environments (ME1 and ME2; A) and single environments included in ME2 (experiments from Peru and Uganda; B). Allelic effects of parental haplotypes to the observed variation explained by the significant QTL on LG 15 (C)

**Table 1.** Summary of QTL analysis results from a multi-environment testing (MET) experiment across 18 environments of Peru, Ghana and Uganda

| QTL | LG | Experiment | Posn (cM) | LSI (cM) | USI (cM) | p-Values | $h^2$ |
| --- | --- | --- | --- | --- | --- | --- | --- |
| 1 | 3 | Ica16C | 3.10 | 0.00 | 37.44 | 2.00E-05 | 22.1 |
| 2 | 13 | Ica16C | 90.20 | 73.91 | 138 | 5.08E-04 | 10.9 |
| 3 | 13 | Ica17C | 106.20 | 87.35 | 130.30 | 1.51E-04 | 15.9 |
| 4 | 15 | Ica16C | 4.90 | 0.00 | 36.02 | 4.16E-05 | 11.5 |
| 5 | 15 | Ica16D | 4.90 | 0.00 | 8.02 | 4.69E-05 | 18.2 |
| 6 | 15 | Puc16 | 6.26 | 0.00 | 43.10 | 1.12E-04 | 14.3 |
| 7 | 15 | Nam16 | 151.18 | 97.15 | 159.00 | 5.87E-05 | 13.2 |
| **8** | **15** | **ME2** | **4.19** | **0.00** | **36.02** | **8.16E-05** | **16.7** |

QTL=quantitative trait loci, LG=linkage group, posn=position, LSI=lower support interval, USI=Upper support interval, $h^2$ =heritability of the QTL, i.e. % variation explained by QTL.

##### Simulated data analysis for misclassification and missingness

Simulations showed that the QTL on LG 15 previously identified by Pereira et al. (2019), which also explained most of the phenotypic variance for rytha (*h*^*2*^ = 20%), had its detection severely reduced as permuted individual proportions increased (Table 2). While this particular QTL was detected as much as 87.5% and 61.0% for 10% and 20% permuted data, respectively, only a 25% detection rate was observed for 30% permuted data. The noise generated by permutation was more prone to detection reduction when compared to analyses involving an increasing proportion of missing data. From 99.0% to 26.0% of detection rate was observed when 10% to 50% of data was missing. For the remaining minor QTL (*h*^*2*^ = 8~11%) previously identified (on LGs 8 and 13), detection rate was consistently low (<22.0%) even for 10% of missing data. On average, logarithm of p-values (LOP) was reduced from 4.49 to 1.35 for 10% to 50% permuted individuals, and from 5.00 to 2.61 for 10% to 50% missing data (**Supplementary Figure 3 and 4**). Regarding the simple trait, FC, two highly significant QTL had been reported (Gemenet et al., 2019; submitted), on LG 3 and LG12. Using simulated data for QTL analysis, the QTL on LG 3 (*h*^2^ = 54%) was consistently detected (>93.5%) regardless of the permutation or missing data proportions, while the QTL on LG 15 (*h*^2^ = 29%) had its detection reduced to 72.0% at 50% of permutation rate, where average LOP went down to 3.54 (**Supplementary Figure 5**). For missing data proportions, LOP was still consistently high (**Supplementary Figure 6**). Due to sampling error and lack of a genome-wide type-I error control, the number of false positives (putative QTL outside the support intervals of QTL previously reported) increased slightly as the proportion of missing data also increased in comparison to permuted data (Table 3).

**Table 2.**
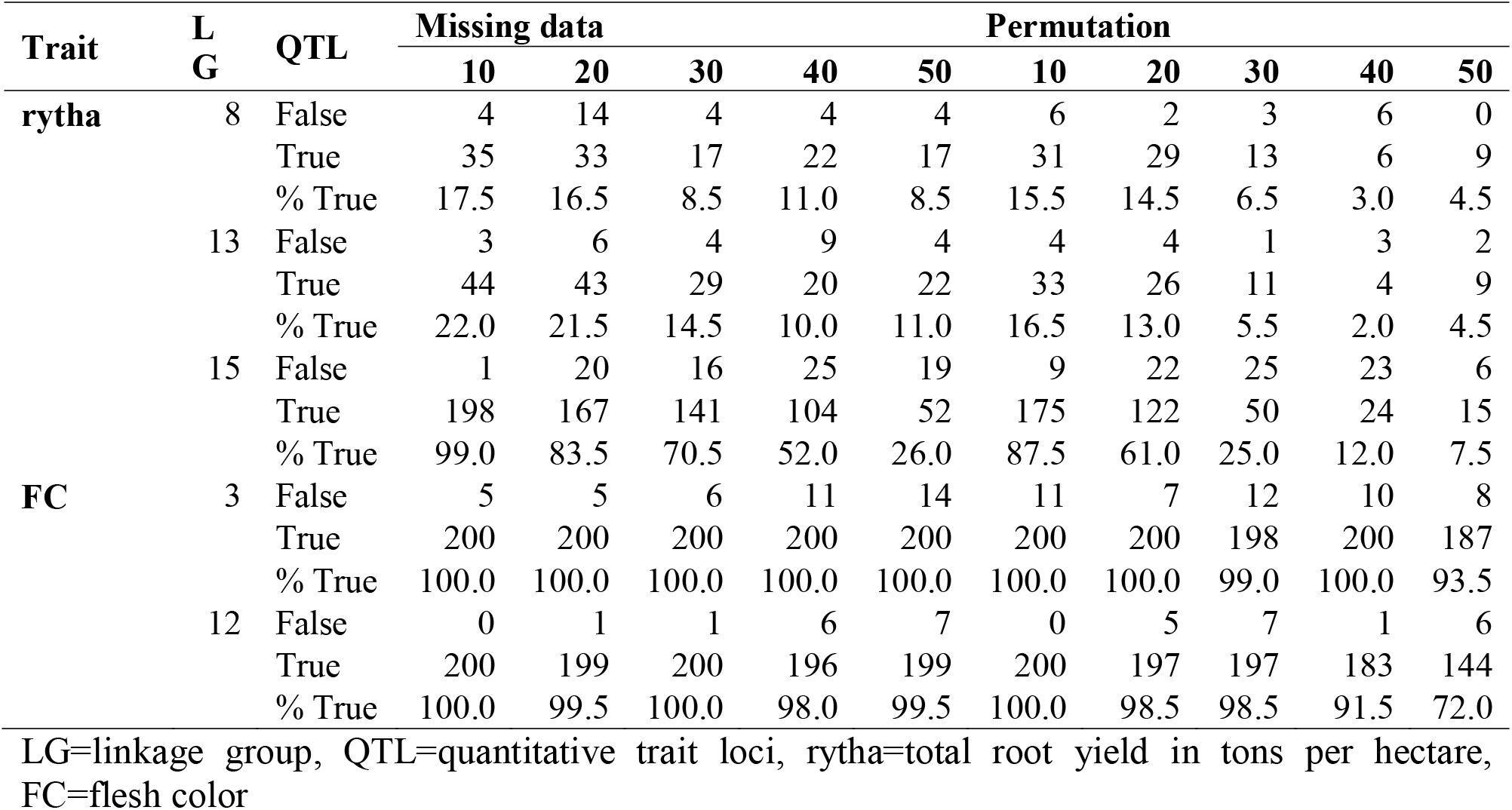
Number of putative QTL in different proportions of missing data and permuted individuals. Each proportion was simulated 200 times. A QTL was considered ‘true’ if located within support intervals of previously identified QTL, and ‘false’ otherwise.

**Table 3.** Number of new putative QTL (regarded as false positives) per linkage group detected in different proportions of missing data and permuted individuals. Each proportion was simulated 200 times.

| Trait | Simulation | Prop (%) | Linkage groups |  |  |  |  |  |  |  |  |  |  |  |  |  |  | Total |
| --- | --- | --- | --- | --- | --- | --- | --- | --- | --- | --- | --- | --- | --- | --- | --- | --- | --- | --- |
|  |  |  | 1 | 2 | 3 | 4 | 5 | 6 | 7 | 8 | 9 | 10 | 11 | 12 | 13 | 14 | 15 |  |
| rytha | Missing | 10 | 72 | 1 | 32 | 3 | 5 | 0 | 2 | 4 | 0 | 0 | 10 | 10 | 3 | 1 | 1 | 144 |
|  |  | 20 | 59 | 3 | 50 | 7 | 2 | 2 | 1 | 14 | 3 | 0 | 18 | 17 | 6 | 13 | 20 | 215 |
|  |  | 30 | 56 | 8 | 39 | 11 | 5 | 6 | 5 | 4 | 2 | 0 | 12 | 24 | 4 | 6 | 16 | 198 |
|  |  | 40 | 52 | 5 | 37 | 15 | 4 | 1 | 11 | 4 | 9 | 3 | 16 | 23 | 9 | 6 | 25 | 220 |
|  |  | 50 | 49 | 8 | 27 | 18 | 1 | 7 | 14 | 4 | 16 | 2 | 18 | 20 | 4 | 11 | 19 | 218 |
|  | Permutation | 10 | 62 | 2 | 34 | 5 | 5 | 1 | 2 | 6 | 0 | 1 | 14 | 16 | 4 | 3 | 9 | 164 |
|  |  | 20 | 42 | 1 | 24 | 7 | 3 | 3 | 3 | 2 | 3 | 1 | 19 | 17 | 4 | 9 | 22 | 160 |
|  |  | 30 | 31 | 5 | 20 | 5 | 4 | 4 | 2 | 3 | 3 | 2 | 12 | 16 | 1 | 2 | 25 | 135 |
|  |  | 40 | 21 | 2 | 17 | 7 | 4 | 5 | 3 | 6 | 7 | 4 | 7 | 7 | 3 | 7 | 23 | 123 |
|  |  | 50 | 13 | 5 | 16 | 6 | 3 | 6 | 8 | 0 | 2 | 6 | 10 | 14 | 2 | 11 | 6 | 108 |
| FC | Missing | 10 | 7 | 2 | 5 | 23 | 0 | 1 | 1 | 0 | 0 | 1 | 0 | 0 | 2 | 6 | 4 | 52 |
|  |  | 20 | 13 | 11 | 5 | 19 | 0 | 0 | 3 | 1 | 2 | 6 | 0 | 1 | 9 | 10 | 1 | 81 |
|  |  | 30 | 11 | 9 | 6 | 15 | 5 | 2 | 7 | 7 | 4 | 2 | 0 | 1 | 6 | 15 | 8 | 98 |
|  |  | 40 | 16 | 19 | 11 | 29 | 7 | 4 | 13 | 4 | 6 | 8 | 3 | 6 | 17 | 16 | 11 | 170 |
|  |  | 50 | 13 | 11 | 14 | 31 | 13 | 5 | 11 | 5 | 11 | 8 | 5 | 7 | 14 | 11 | 14 | 173 |
|  | Permutation | 10 | 10 | 10 | 2 | 11 | 2 | 1 | 3 | 2 | 2 | 12 | 1 | 0 | 4 | 12 | 4 | 76 |
|  |  | 20 | 12 | 17 | 3 | 19 | 7 | 3 | 4 | 3 | 3 | 8 | 1 | 5 | 10 | 11 | 6 | 112 |
|  |  | 30 | 10 | 8 | 12 | 14 | 11 | 3 | 7 | 2 | 5 | 5 | 7 | 7 | 12 | 5 | 7 | 115 |
|  |  | 40 | 11 | 8 | 8 | 12 | 6 | 4 | 7 | 4 | 7 | 7 | 4 | 1 | 18 | 5 | 4 | 106 |
|  |  | 50 | 16 | 11 | 8 | 10 | 4 | 3 | 6 | 3 | 7 | 3 | 6 | 6 | 10 | 8 | 8 | 109 |

Prop=proportion, rytha=total root yield per hectare, FC=flesh color.

## Discussion

### Case Study 1: More than 20% pedigree error likely to affect future predictions based on the MDP population negatively

Experimental noise is detrimental to studies seeking to combine phenotypic and genomic data such as QTL analysis, genome-wide association mapping and genomic selection. In Case Study 1, we found 27.7% error due to mislabeling from *in vitro*, screenhouse and field, and 22.7% error for mislabeling between and within families. The difference between the two errors is that the former just indicates whether a genotype retains the same label from *in vitro*, screen house and field regardless of family assignment, whereas the former looks at clustering based on family assignment. Effects of genotype mislabeling have been reported in humans, animals and plants. Buyske et al. (2009) showed that to retain the same power for marker-trait association in humans, a 39-fold more sample size was required if the mislabeling error was 5%. Long et al. (1990) observed that a 20% error in pedigree labeling resulted in 9.3, 3.2 and 12.4% reduction in genetic gain when using phenotypic BLUPs, for litter size, backfat and average daily gain, respectively, in pigs. This implies that the degree of sensitivity to pedigree errors are also influenced by trait architecture. Using an F_1_ population data previously analyzed for rytha (Pereira et al., 2019) and FC (Gemenet et al., 2019; submitted) in sweetpotato, we noticed that QTL detection was in fact more severely impacted for traits with lower heritability, like rytha, compared to high heritability traits like FC, when permuted data was simulated (Table 2 and **Supplementary Table 4**).

Mislabeling between families is also expected to have a negative effect on predictions especially in breeding programs where full-sib and half-sib family means are used in selection. In pigs, it was shown that 20% pedigree errors reduced genetic gain by 7.0, 2.5 and 7.5% in litter size, backfat and average daily gain, respectively, when using family means for selection (Long et al., 1990). In sweetpotato at CIP, most breeding programs are now adopting population hybrid breeding schemes which rely on progeny testing for selection. The current data does not allow estimation of the reduction in genetic gain expected when using either breeding values and family means for selection since we have not genotyped the whole population yet and also our experiment is not designed in a case-control manner. In plants, several prediction models were tried out to identify those that are relatively tolerant to pedigree errors using sugar beet. The study by Biscarini et al. (2016) indicated that local classification methods such as K-nearest neighbor and random forest tolerated the pedigree noise better compared to methods using global data properties. Knowing the estimated pedigree errors in the current Case Study 1 is important as it will allow to explore such tolerant methods for breeding value prediction in future analyses and decisioning.

### Case Study 2: Lower genetic correlations and lack of significant QTL in some mega-environments indicate a level of phenotype misclassification in the mapping population

Differential QTL expression in relation to environmental variables is expected in MET analyses especially for complex quantitative traits (Boer et al., 2007). In our Case Study 2, we used mixed models to account for GE interactions as well as the associated genetic correlation structures and extended these to QTL mapping by matching the phenotypes to the respective genotypes as covariates. Following MET analysis, only one of the QTL identified in SEs of ME2 was stable across the ME2. The QTL on LG 15 at position 4.19 cM can therefore be classified as a constitutive QTL for rytha in sweetpotato in the current genetic background whereas the other QTL on LG 3, 13 and LG 15 position 151.18 cM as adaptive QTL for those specific environments (Vargas et al., 2006). The only QTL of ME2 was mapped before in a combined analysis of all environments in Peru and possible candidate genes underlying this QTL are described in Pereira et al. (2019). Based on phenotypic data, the environments from Peru were all correlated possibly due to better data quality and the availability of field maps for spatial adjustments using rows and columns, when compared to the environments in Uganda and Ghana. These improved correlations in Peru were also extended to the QTL results where most of the significant QTL in SEs were identified in environments from Peru, with only one QTL being identified in one environment of Uganda. These results therefore confirm the findings of previous studies that spatial adjustment improves genetic correlations among environments (Lado et al., 2013; Elias et al., 2018; Ward et al., 2019). Although rytha, like many yield traits is quantitative and prone to GE interaction and QTL x E interaction (Boer et al., 2007; van Eeuwijk et al., 2010), we did not observe different QTL for the two MEs, rather, there was no significant QTL for ME1. Additionally, significant adaptive QTL were mainly identified for the environments from Peru. We therefore hypothesize that since population development and DNA extraction was carried out in Peru and no QC/QA was carried out for trials in Uganda or Ghana after shipping of *in vitro* genotypes, phenotype misclassification may have occurred in some environments. This would lead to phenotypes from some of the environments not matching entirely with the genotypic data, consequently resulting in lack of association between the trait and the markers, as demonstrated by simulations. This hypothesis is also supported by the fact that zero correlation was observed between some environments in Uganda and Ghana with the rest of the environments in the same country, at the single environment (SE) analysis step. Looking at the MEs defined at the second analytical stage, the genetic correlations among the environments in ME1 were lower than those observed among environments of ME2, even though ME2 contained environments from both Peru and Uganda while ME1 contained environments only from Ghana. Consequently, no QTL could be identified for ME1. Although a percentage of this can be attributed to GE interaction, we assume that the lack of correlation from one environment with the next could be a result of a certain degree of misclassification. Given that QTL detection in complex traits like rytha is difficult due to low trait heritability, we confirmed presence of misclassification by analyzing QTL in ME1 for a simple trait, flesh color, and the known high effect QTL already reported in (Gemenet et al., 2009; submitted) could not be captured either (data not shown). The QTL for ME2 could only explain 16.7% of the observed variation which is expected for complex traits like rytha as only QTL with relatively higher effect can be captured in QTL tagging based on biparental populations, thereby leading to the concept of missing heritability (Crossa, 2012).

### Phenotype misclassification affects QTL detection more than missingness, and the magnitude of effects is trait specific

Our simulation results showed that misclassification leads not only to decreased detection power, but also to QTL contribution underestimation. We also showed that especially for complex traits, phenotype misclassification had more detrimental effects on QTL detection when compared to missingness in data. Additionally, we found that although both misclassification and missingness affected QTL detection in simple traits as proportions increased, the reduction in QTL detection was much lower compared to quantitative traits. Missingness is a well-documented problem especially in human genetics where some phenotypes are difficult to measure in large enough populations for effective marker-trait association studies (Jiang et al. 2018). Missingness can lead to both type-I and type-II error in analysis and several methods have been proposed to mitigate against (Hormozdiari et al. 2016; Jiang et al. 2018; Chen et al. 2018). Plant breeding datasets are always unbalanced due to missingness. Similar to our results, Galli et al. (2018) showed that although missingness slightly reduced predictive ability as proportions of missing data increased, the selected fraction was not much affected in genomic prediction of maize hybrids. Contrastingly, misclassification has more dramatic effects even for simple traits. for example, even though the simulation data used in our study based on good quality phenotypic data from Peru indicates that misclassifications resulted in lower effects on QTL detection in simple traits, analyzing flesh color, a simple trait from ME1 above made up of environments from Ghana in which most misclassification and pedigree errors are suspected did not capture the high effect QTL already reported for the same trait in several other populations and environmental backgrounds. Therefore, addressing misclassification and pedigree errors requires proper attention in breeding trials to enhance increased genetic gains.

### Implications for the sweetpotato breeding programs

We have demonstrated the estimated level of pedigree error due to genotype misclassification by mislabeling and also demonstrated the likely consequences of such errors when combining the phenotypes with the respective genotypes using two case studies, and simulated data. Since breeding programs are moving more and more into genomic selection, genetic gain from such breeding activities will depend on the accuracy of predicting untested genotypes. Several methods have been proposed to help improve such prediction accuracy and could be adopted for sweetpotato breeding. Modeling of GE interaction and spatial adjustment using mixed models is one way to cater for environmental heterogeneity and improve such prediction accuracy (Piepho, 1998; Burgueno et al., 2011; de los Campos et al., 2009; Crossa et al., 2010, 2011; Bernal-Vasquez et al., 2014). In this study we have observed that genetic fidelity due to proper labeling combined with spatial adjustment where necessary, improved the genetic signal for tagging QTL. Ability to tag QTL is important because linkage disequilibrium between markers and QTL is important in improving prediction accuracy in genomic selection (Nakaya and Isobe, 2012; Spindel et al., 2016). Additionally, use of multi-trait, multi-environment prediction has been shown to improve prediction accuracy (Covarrubias-Pazaran et al., 2018; Sun et al., 2017; Mitchel et al., 2019) and genetic gain from genomics-assisted breeding. Such multi-trait analyses, also known as multivariate analyses, take advantage of the genetic correlations between simple secondary traits and complex yield traits included in both training and prediction populations to improve prediction accuracy of the complex trait.

In case of misclassification and pedigree errors, a few statistical approaches have been explored to improve prediction accuracy such as using less sensitive prediction methods (Biscarini et al., 2016) and the use of realized relationship matrices to correct pedigree errors (Munoz et al., 2013). These can be adopted in sweetpotato especially in the case of the MDP population described in Case Study 1 to improve prediction accuracy of future studies. However, the use of improved statistical analytic methods is a reactionary approach to improving prediction accuracy and marker-trait associations, and its benefits may be limited depending on how much human error is present in a given trial. To better take advantage of the advances in genomics-assisted breeding, sweetpotato breeding would make faster genetic gains from adopting improved breeding process and plot management practices to avoid both pedigree errors due to genotype misclassification and experimental errors. This would require putting in place and applying next generation data management and analytical decision support tools for participating sweetpotato breeding programs (Rathore et al., 2018). We propose that the sweetpotato breeding process be mapped out and documented in each breeding program. Additionally, standard operating procedures (SOPs) should be documented and implemented at each stage of the breeding process such as: crossing and seed inventory, experimental designs and trial establishment, phenotyping and digitalized data capture, standard trait ontologies, data checks and quality metrics, meta-data recording, sample tracking and genotyping workflows, marker-assisted selection and genomic selection. Additionally, barcoding and QC/QA of breeding and trialing populations should be adopted and applied routinely.

## Data Availability

The SNP data used in Case study 1 and the phenotypic data used in Case study 2 are submitted together with this manuscript, as Supplementary Table 1 and Supplementary Table 2, respectively. The genetic linkage map used in QTL mapping can be interactively accessed at (https://gt4sp-genetic-map.shinyapps.io/bt_map/).

## Author Contributions

DG, MK, RS, OU, JS, EC, WG, BY, CY, RM planned and carried out field and laboratory experiments, DG, BB, GP carried out data analysis, DG wrote the manuscript. All authors read and approved the manuscript.

## Funding

The research was supported by a Bill & Melinda Gates Foundation grant (Grant number OPP1052983), as part of the Consultative Group on International Agricultural Research (CGIAR)-Research Program on Roots, Tubers and Bananas (RTB) which is supported by CGIAR Fund Donors (http://www.cgiar.org/about-us/our-funders/).

## Acknowledgements

The authors wish to acknowledge the scientific and technical teams at CIP-Peru, CIP-Uganda and CIP-Ghana for carrying out field experiments and the team at CIP-Kenya for carrying out laboratory analyses. Additionally, the authors acknowledge the Genomic Tools for Sweetpotato Improvement project (GT4SP) members for indirect contribution towards the manuscript.

## Conflict of Interest

The authors declare no conflict of interest.

---

**Supplementary Figure 1**. A zoomed out phylogenetic tree (from Figure 1), showing the clustering of some genotypes selected from *in vitro* (red with prefix I), screenhouse (black with prefix S) and field (blue with prefix F). The majority of genotypes are consistently clustered. However, some of them are not e.g. I28 which is clustering differently from S28 and F28.

**Supplementary Figure 2**. A map showing the experimental locations in Peru, Ghana and Uganda used in the evaluation of a biparental mapping population.

**Supplementary Figure 3**. QTL mapping for increasing proportion of permuted individuals (10, 20, 30, 40 and 50%) from the non-permuted, original data (0%) for rytha (Pereira et al. 2019). Colored lines are their respective LOP average for 200 simulations each.

**Supplementary Figure 4.** QTL mapping for increasing proportion of missing data (10, 20, 30, 40 and 50%) from the non-missing, original data (0%) for rytha (Pereira et al. 2019). Colored lines are their respective LOP average for 200 simulations each.

**Supplementary Figure 5.** QTL mapping for increasing proportion of permuted individuals (10, 20, 30, 40 and 50%) from the non-permuted, original data (0%) for FC (Gemenet et al. 2019). Colored lines are their respective LOP average for 200 simulations each.

**Supplementary Figure 6.** QTL mapping for increasing proportion of missing data (10, 20, 30, 40 and 50%) from the non-missing, original data (0%) for FC (Gemenet et al. 2019). Colored lines are their respective LOP average for 200 simulations each.

**Supplementary Table 1.** DArTseq single nucleotide polymorphism markers used in Case Study 1

**Supplementary Table 2.** Genotype list and family assignments of genotypes used in Case Study 1

**Supplementary Table 3.** Raw data from field experiments in Peru, Ghana and Uganda measured on a biparental mapping population used in Case Study 2.

