## Supplementary figures and images for "When a phenotype is not the genotype: Implications of phenotype misclassification and pedigree errors in genomics-assisted breeding of sweetpotato [*Ipomoea batatas* (L.) Lam.]"

### Supplementary Figure 1

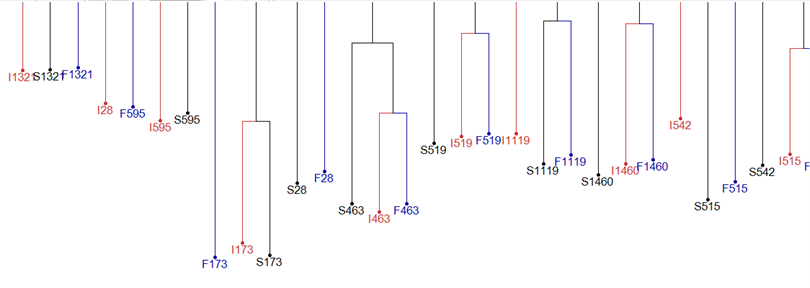

### Supplementary Figure 2

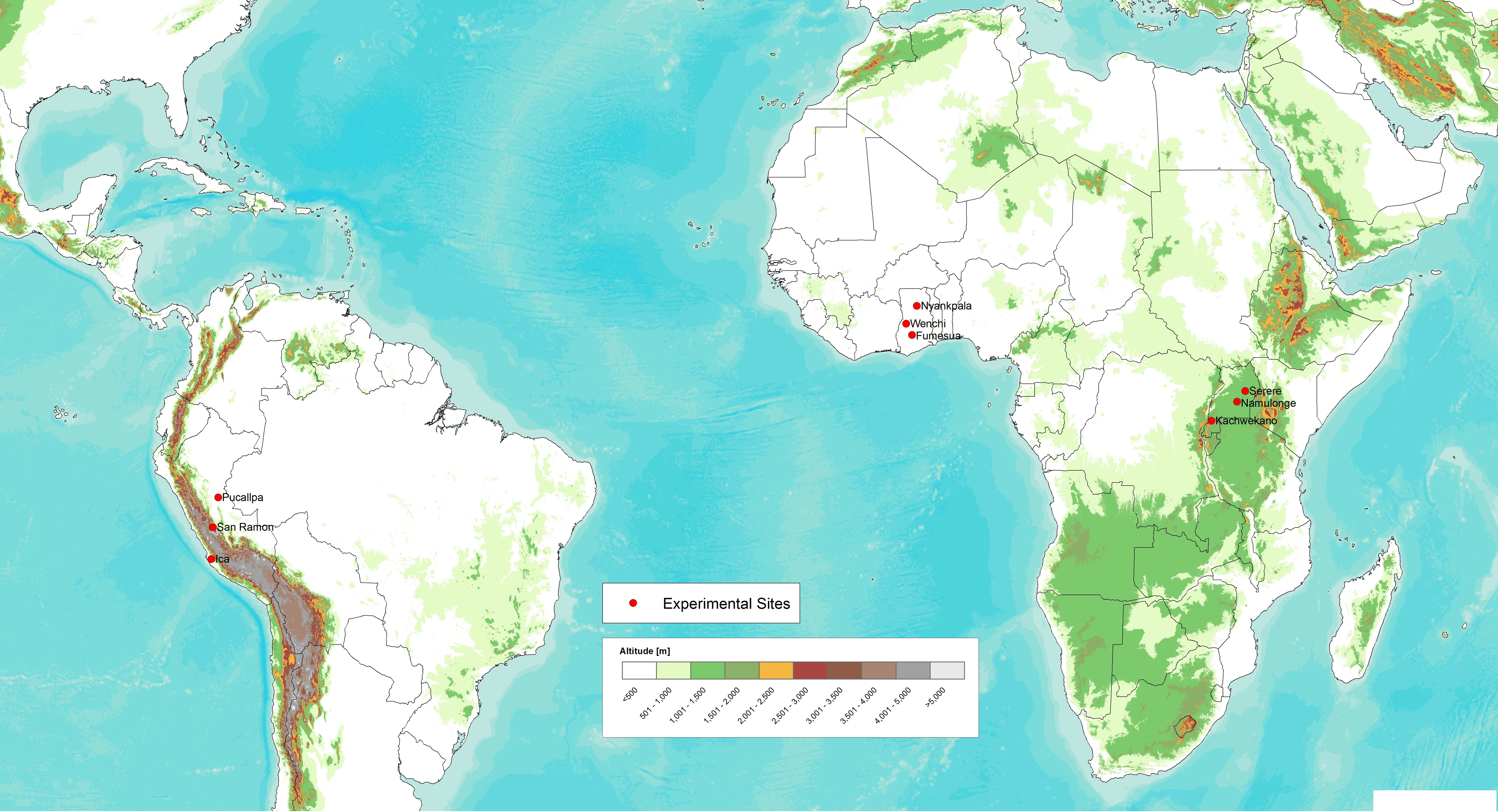

### Supplementary Figure 3

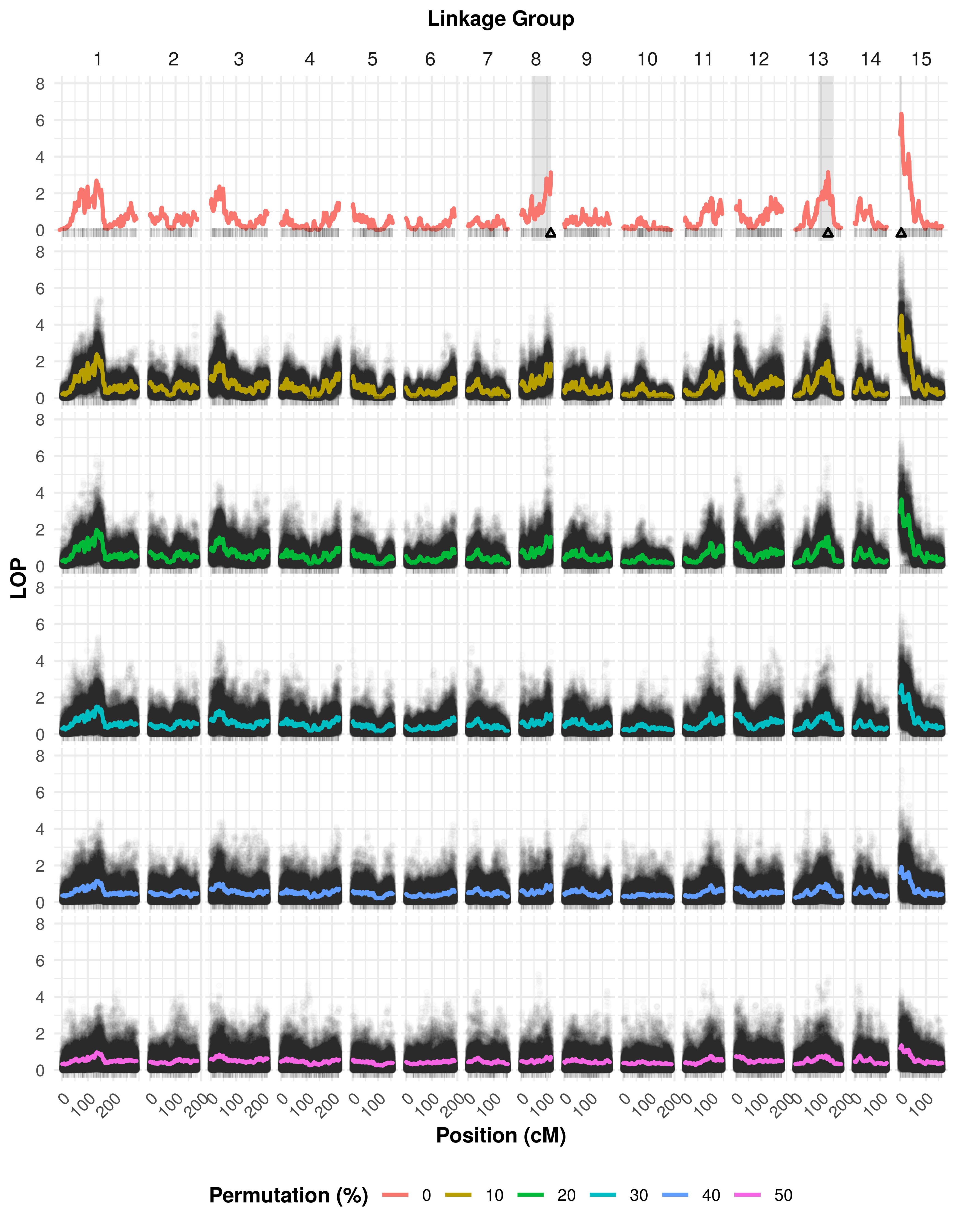

### Supplementary Figure 4

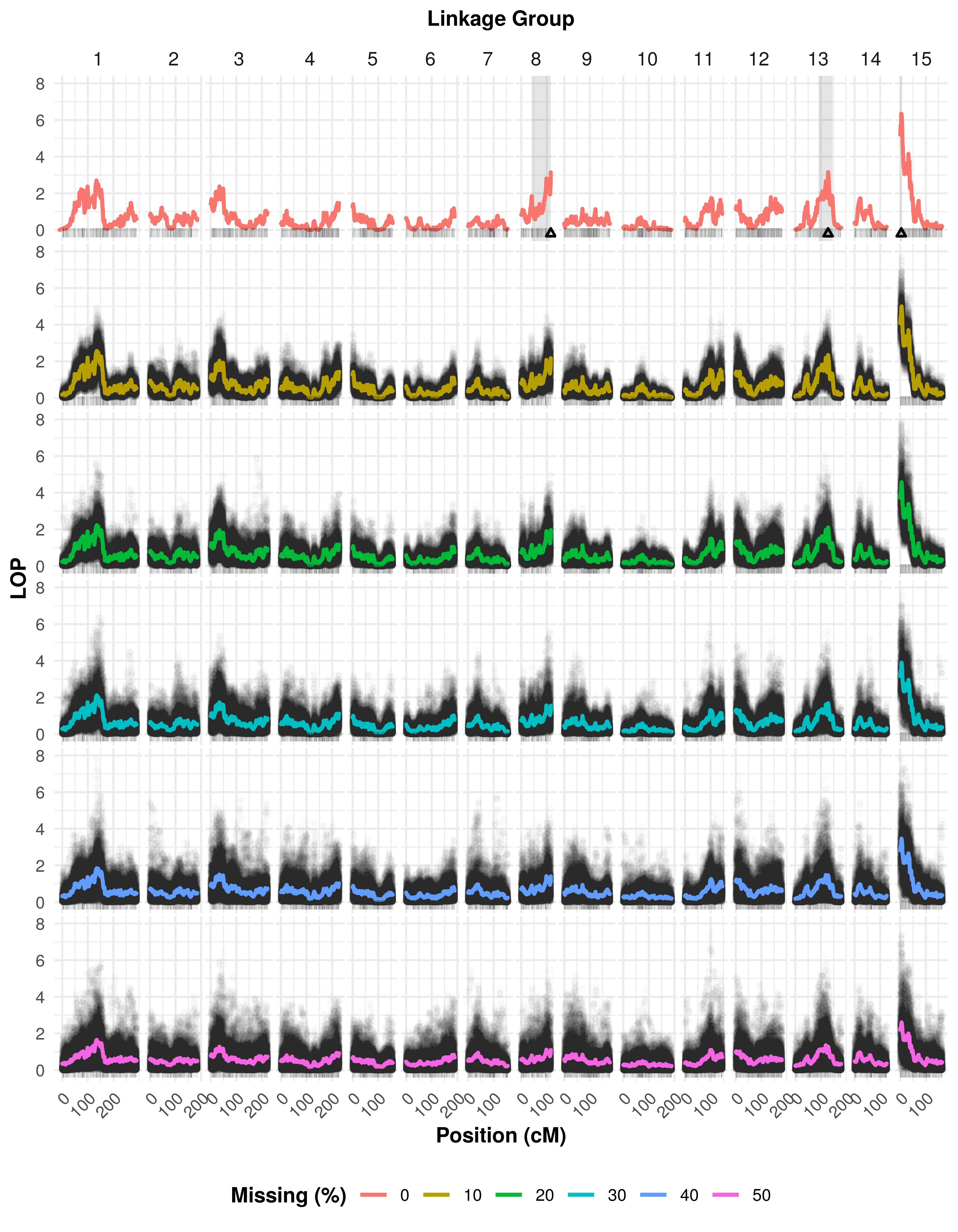

### Supplementary Figure 5

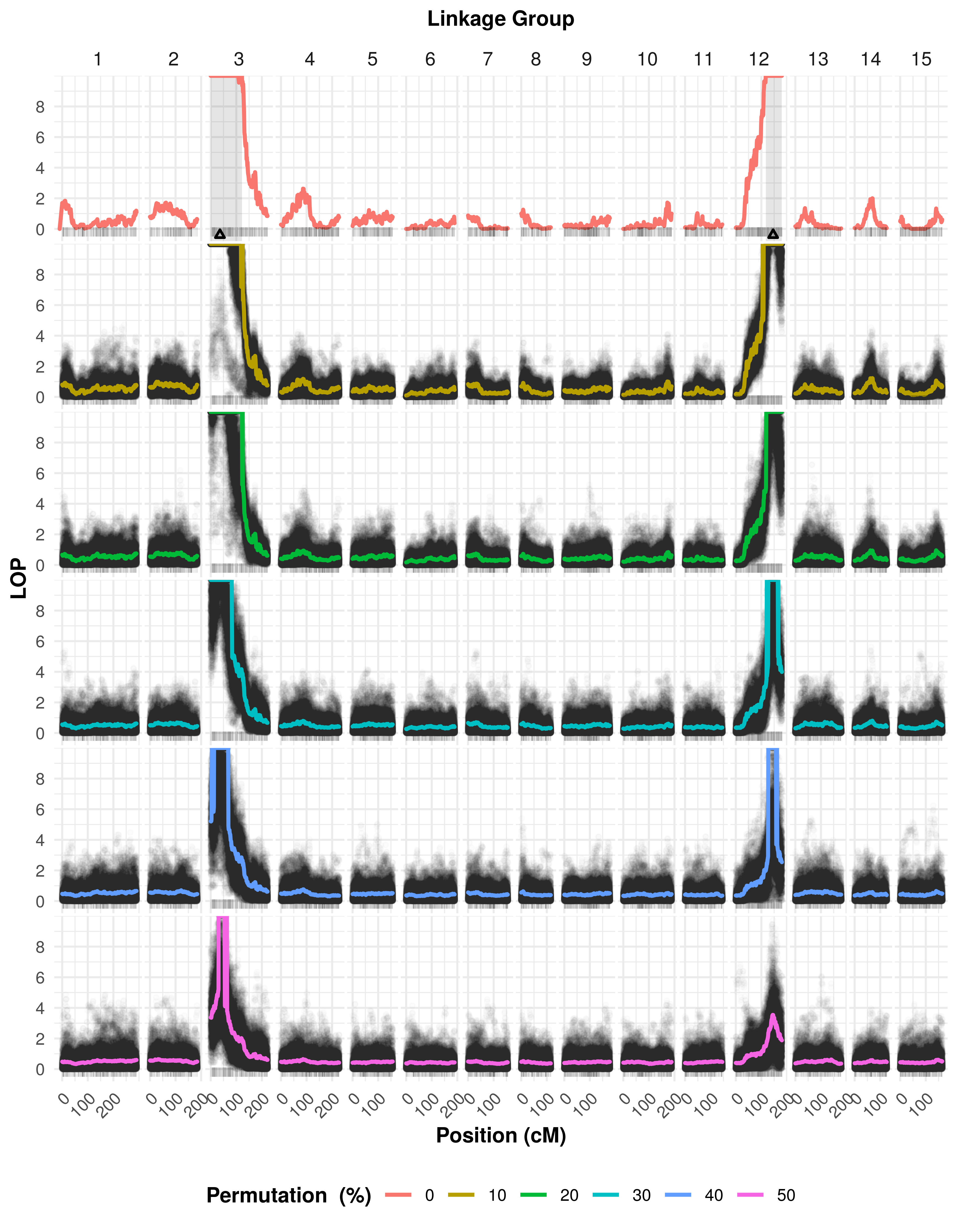

### Supplementary Figure 6

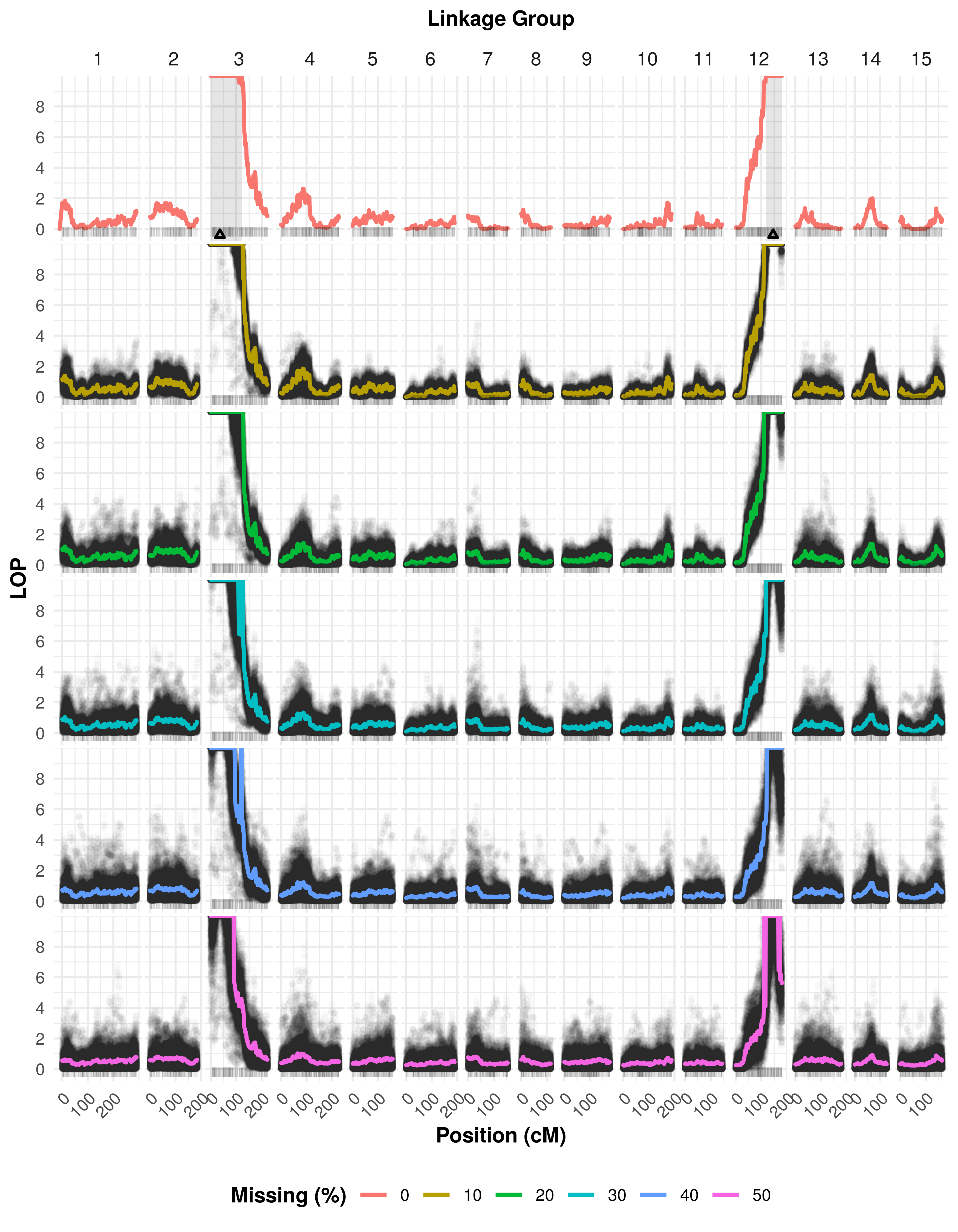
